# Transcription dynamics and regulation of heat shock protein genes during stress and development in the estuarine cnidarian *Nematostella vectensis*

**DOI:** 10.1101/2025.10.15.682611

**Authors:** Janki A. Bhalodi, Joachim M. Surm, Adam M. Reitzel

## Abstract

Heat shock proteins (HSPs) are conserved molecular chaperones that function in protecting cells from proteotoxic conditions. Most eukaryotes have multiple HSP genes that encode for proteins localizing in the cytoplasm, endoplasmic reticulum (ER), and mitochondria. In marine invertebrates of the phylum Cnidaria, where HSPs are often used as biomarkers of environmental stress, little is known about how particular HSPs vary in copy number, expression, inducibility, and regulation within a species. In this study, we characterized the diversity of the full repertoire of HSP70 and HSP90 genes in an emerging model cnidarian, *Nematostella vectensis*. We identified five distinct HSP70 and three HSP90 genes, with at least one homolog from each family belonging to the three primary clades based on subcellular localization. Although transcription of none of these HSPs was significantly induced by a temperature change of 10°C, two cytosolic HSP70s and one cytosolic HSP90 were significantly upregulated with a 20°C increase in temperature. Most HSPs followed a similar pattern of expression across various developmental stages of *N. vectensis*, with an increase in expression during the early larval stage followed by a decrease during the juvenile stage. HSPs showed evidence for differential expression in particular cell types, where multiple cytosolic and ER HSPs were highly expressed in neuronal and cnidocyte cell types. Moreover, there were differences in the abundance and sequences of regulatory heat shock element motifs in the putative promoters of *N. vectensis* HSPs, providing a potential mechanistic basis of functional diversification in response to temperature and development. By characterizing the expression of all HSP70 and HSP90 genes in this cnidarian, we reveal distinct roles of these core genes in the proteostasis response, providing a foundation for future functional studies into the diverse contributions of HSPs to cnidarian life cycle and stress resilience.

## 1. Introduction

Organisms across the tree of life are exposed to environmental stressors that can disrupt cellular homeostasis. To cope with these challenges, species have evolved diverse pathways for responding to environmental stressors. One particularly conserved network of molecular chaperones is the heat shock protein (HSP) family, which facilitates protein folding, prevents aggregation, and helps reestablish protein homeostasis within the cell [1–4]. During cellular stress, the transcription of HSP genes is primarily regulated by the DNA-binding heat shock factor (HSF), which recognizes heat shock element (HSE) motifs consisting of three or more inverted repeats of the 5’- nGAAn-3’ pentameric DNA sequence [5]. HSPs are classified into distinct protein families based on molecular weight, including large HSPs, HSP90, HSP70, HSP60, HSP40, and small HSPs; members of the HSP70 and HSP90 families play central roles in cellular stress responses [6, 7] and have been widely used as biomarkers of stress across species [1]. HSP70 and HSP90 proteins also play critical roles in development, contributing to cellular differentiation [8], cell cycle signaling pathways [9], and regulation of apoptosis [8]. Particular members of the HSP70 and HSP90 families, often referred to as “constitutive” or “heat shock cognate”, have essential roles in cellular housekeeping such as protein folding and unfolding, transport, and the degradation of misfolded or aggregated proteins [1, 2, 6, 7]. In eukaryotes, the HSP70 and HSP90 families diversified into three major phylogenetic clades corresponding to distinct cellular localizations [10]. Within these clades, the number of HSP genes as well as their expression varies extensively [11], pointing to functional differences between HSPs localizing in different cellular organelles. In a highly duplicated family such as the HSPs, such variation can result from subfunctionalization or neofunctionalization of paralogs [12, 13].

Although HSPs are present across nearly all domains of life, their copy number, expression, inducibility, and regulation vary among species and environments, suggesting lineage-specific adaptations to distinct ecological niches and physiological demands [14–18]. While HSP diversity and regulation have been characterized broadly across other taxa, much less is known about how these dynamics manifest in species in the phylum Cnidaria (e.g., corals, sea anemone, jellyfish, hydras), where HSPs are often used as biomarkers of cellular stress. HSPs have been studied in a variety of cnidarian species where their expression is generally correlated with responses to stress across species [19]. However, many of these studies tend to identify the genes under study as ‘HSP70’ or ‘HSP90’, which overlooks the complexity of these chaperone families. For instance, previous studies in the corals *Acropora digitifera* and *Acropora millepora* examined HSP70 and HSP90 gene expression under acidified seawater and bacterial challenges, respectively [20, 21], but did not capture HSPs from the full range of subcellular localization-based phylogenetic clades.

Previous studies in various cnidarian species have shown variation in the diversity and expression of HSPs within and between species. For example, the intertidal sea anemone *Anthopleura elegantissima* exhibited population-specific patterns of HSP70 expression that correlated with thermal microhabitats, highlighting environmentally driven divergence in inducibility and baseline expression even within a single species [15]. Similarly, while *Hydra attenuata* and *Hydra magnipapillata* exhibit both heat shock protein induction and thermotolerance, these responses are absent in the closely related *Hydra oligactis*, highlighting species-specific variation in stress response capacity among cnidarians [22, 23]. In other cnidarians, HSPs have been linked to developmental functions. For instance, in *Hydractinia echinata,* an HSP70 protein was shown to function in cell proliferation, anti-apoptosis, and axial patterning [24], while in the coral *Acropora millepora*, HSP expression changes were shown to accompany metamorphosis [25]. Given that many metazoan HSPs are implicated in developmental processes, further study of cnidarian HSPs is needed to better understand their roles outside of stress responses. Collectively, these findings underscore that even highly conserved chaperone families can exhibit substantial regulatory and functional divergence depending on ecological and developmental context. Consequently, resolving the diversity and expression patterns of all HSPs at the species level is essential for understanding the specific functional roles of each HSP gene and the overall physiological variability in stress responses across taxa.

*Nematostella vectensis*, a burrowing cnidarian sea anemone, inhabits highly variable estuarine environments where temperature, salinity, and oxygen levels can shift dramatically over short time scales [26–28]. *N. vectensis* has a broad geographic distribution, ranging from the Atlantic and Pacific coasts of North America to parts of the United Kingdom [29, 30]. This species lives in environments where it experiences temperatures very close to its physiological limits [27]. Evidence suggests local adaptation to temperature in geographically distinct populations of *N. vectensis,* with lower-latitude populations exhibiting reduced HSP induction under heat stress [31] and higher growth rates at the same temperature [27]. A genomic survey identified six HSP70 genes, three HSP90 genes, and an HSF1 homolog in the *N. vectensis* genome [32], suggesting conservation of this stress response mechanism in sea anemones. *N. vectensis* upregulates HSPs in response to multiple stressors, including heavy metals [33], heat [34–36], and oxidative stress [37]. Functional differences in protein chaperone function between three *N. vectensis* HSP70 proteins have been explored through heterologous expression in *Saccharomyces cerevisiae*, revealing differing capacities to confer resistance against various stressors [34]. *N. vectensis* HSP70s also interact with unique sets of client proteins and co-chaperones in the absence of stress, which are then modulated upon heat shock [38]. Stress response characterization in *N. vectensis* has largely focused on HSP70 expression, and there is a lack of data on HSP90s mediating any aspect of function, including the cellular stress program. In this study, we characterized the diversity of *N. vectensis* HSP70 and HSP90 genes through phylogenetic comparisons, analysis of their expression during heat stress and development, and identification of regulatory HSE motifs in their promoters. By describing the diversity and expression of each HSP gene, our study provides a thorough description of potential functional variation for these common cellular chaperones in an emerging animal model species.

## 2. Results

### 2.1. Identification and phylogenetic analysis of candidate HSP70 proteins

We identified six HSP70 proteins in the *N. vectensis* genome. Upon further examination of this set, we found a pair of identical proteins that seemed to have arisen from a duplication event on chromosome 4, resulting in two identical HSP70 genes located 6654 bp apart in opposing orientations. Since these genes also had identical putative promoter sequences and there was no way we could differentiate between their expressions, we only included one of these in our study (Hsp70A2). The resulting set of five HSP70s ranged between 640-691 amino acids in length and were predicted to have molecular weights between 69.9-75.12 kDa (Table 1). Subcellular localization predictions revealed that Hsp70A1A, Hsp70A1B, and Hsp70A2 are all predicted to localize in the cytoplasm, while Hsp70B1 and Hsp70C1 are predicted to localize in the endoplasmic reticulum and mitochondria, respectively (Table 1).

**Table 1.** Summary of the HSP70 protein family in *N. vectensis*.

| Gene name | Protein length (aa) | Molecular weight (kDa) | Subcellular localization | Refseq gene ID | Refseq protein ID |
| --- | --- | --- | --- | --- | --- |
| Hsp70A1A | 655 | 71.99 | Cytoplasm | 5508770 | XP_001629343.2 |
| Hsp70A1B | 653 | 71.41 | Cytoplasm | 5501070 | XP_001622423.1 |
| Hsp70A2 | 640 | 69.9 | Cytoplasm | 5516533 | XP_001636593.2 |
| Hsp70B1 | 669 | 73.88 | Endoplasmic reticulum | 5504498 | XP_001625396.1 |
| Hsp70C1 | 691 | 75.12 | Mitochondrion | 5519914 | XP_001639786.3 |

We also investigated the phylogenetic relationships among *N. vectensis* HSP70s and those from other organisms, including annotated genes from humans, flies, and nematodes, to understand how anemone genes are related. All *N. vectensis* HSP70s are grouped with their respective cytosolic, endoplasmic reticulum, and mitochondrial orthologs in other species (Figure 1). NvHsp70A1A and NvHsp70A1B are most closely related to each other and other cnidarian HSP70s; however, NvHsp70A2 segregated into a clade that included fly and worm HSP70s (Figure 1). NvHsp70B1 is grouped with human, insect, and other cnidarian HSP70s in the ER clade (Figure 1). Similarly, NvHsp70C1 shares a common ancestor with coral, hydra, human, and nematode HSP70s, all of which are in the mitochondrial clade (Figure 1).

**Figure 1.**
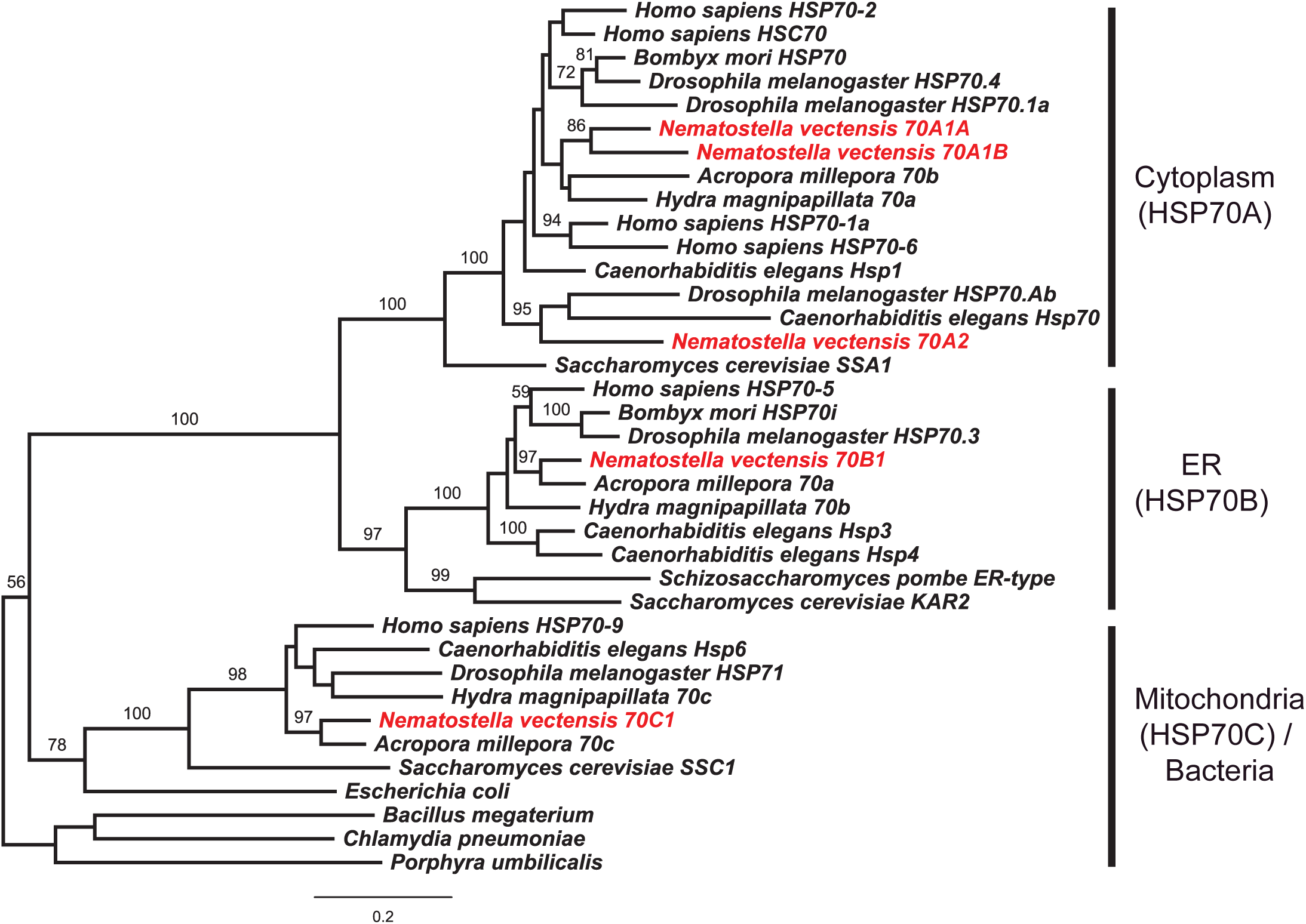
Phylogenetic relationships between HSP70 proteins from *N. vectensis* and other species. Phylogeny was constructed using the maximum likelihood method with 1000 bootstrap replicates. Bootstrap values above 50 are indicated on branches. HSP70 clades are labeled (right) with the subcellular localization of proteins. *N. vectensis* HSPs are colored red.

### 2.2. Identification and phylogenetic analysis of candidate HSP90 proteins

We identified three candidate HSP90s in the *N. vectensis* proteome. These proteins have predicted molecular weights ranging from 83.35-97.8 kDa, which is within the typical range of weights for proteins belonging to the HSP90 family (Table 2). The protein lengths of candidate *N. vectensis* HSP90s range from 727-847 amino acids (Table 2). Hsp90A1 is predicted to localize in the cytoplasm, Hsp90B1 in the endoplasmic reticulum, and NvTRAP in the mitochondria (Table 2).

**Table 2.** Summary of the HSP90 protein family in *N. vectensis*.

| Gene name | Protein length (aa) | Molecular weight (kDa) | Subcellular localization | Refseq gene ID | Refseq protein ID |
| --- | --- | --- | --- | --- | --- |
| Hsp90A1 | 727 | 83.35 | Cytoplasm | 5520770 | XP_001640689.1 |
| Hsp90B1 | 847 | 97.8 | Endoplasmic reticulum | 5517472 | XP_001637407.2 |
| NvTRAP | 751 | 85.54 | Mitochondrion | 5517716 | XP_032218193.2 |

When compared to HSP90 proteins from other species, *N. vectensis* proteins group with their respective orthologs based on their subcellular localization (Figure 2). NvHsp90A1 and NvHsp90B1 both share more recent common ancestors with human, insect, and nematode HSP90s than they do with plant HSP90s (Figure 2). NvTRAP clusters with human TRAP with moderate support and shares a more recent common ancestor with insect TRAPs compared to the nematode TRAP, all of which group in the mitochondrial clade of HSP90s (Figure 2).

**Figure 2.**
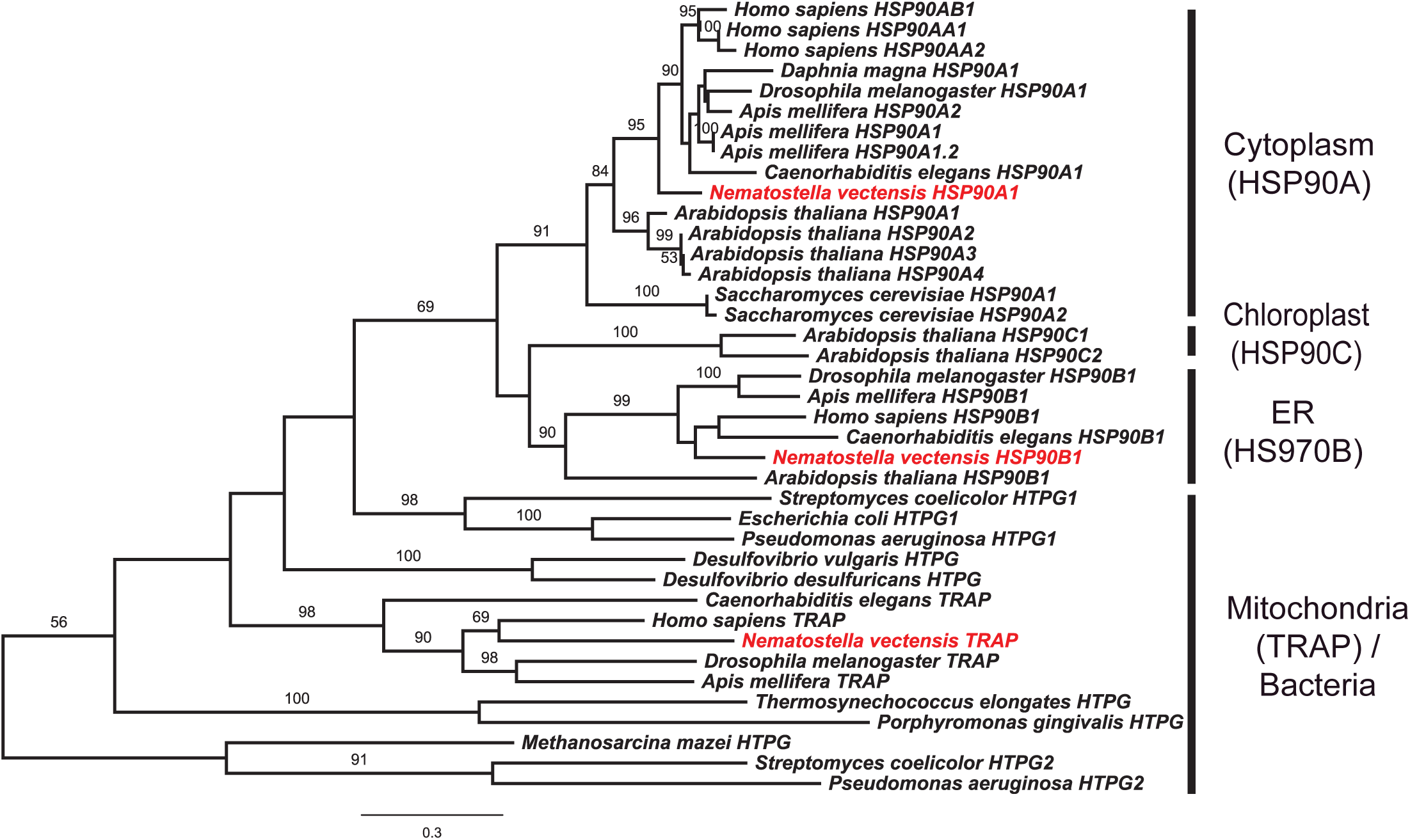
Phylogenetic relationships between HSP90 proteins from *N. vectensis* and other species. Phylogeny was constructed using the maximum likelihood method with 1000 bootstrap replicates. Bootstrap values above 50 are indicated on branches. HSP90 clades are labeled (right) with the subcellular localization of proteins. *N. vectensis* HSPs are colored red.

### 2.3. Expression of HSP70s and HSP90s during acute temperature stress

To determine the expression of each *N. vectensis* HSP gene in response to temperature changes, we tested the inducibilities of each HSP over a range of environmentally relevant temperatures experienced by the anemones in their natural habitats. The transcript counts for each HSP at various temperatures (Figure 3A) were normalized to those at 20°C to visualize the fold change in expression relative to the control expression (Figure 3B). All HSPs were expressed at some level at the control temperature of 20°C, with Hsp70A1B having the highest expression (Figure 3A). None of the HSPs showed a significant change in transcript levels at 10°C or 30°C (Figures 3A and 3B). In contrast, at 40°C, Hsp70A1A, Hsp70A2, Hsp70B1, Hsp90A1, and Hsp90B1 were all upregulated, while Hsp70A1B was downregulated (Figures 3A and 3B). Hsp70C1 and NvTRAP were the only HSPs that did not show a significant change in expression (Figure 3A). Although all other HSPs had significant changes in expression at 40°C, the highest induction was exhibited by Hsp70A2, followed by Hsp90A1 and Hsp70A1A, respectively (Figures 3A and 3B).

**Figure 3.**
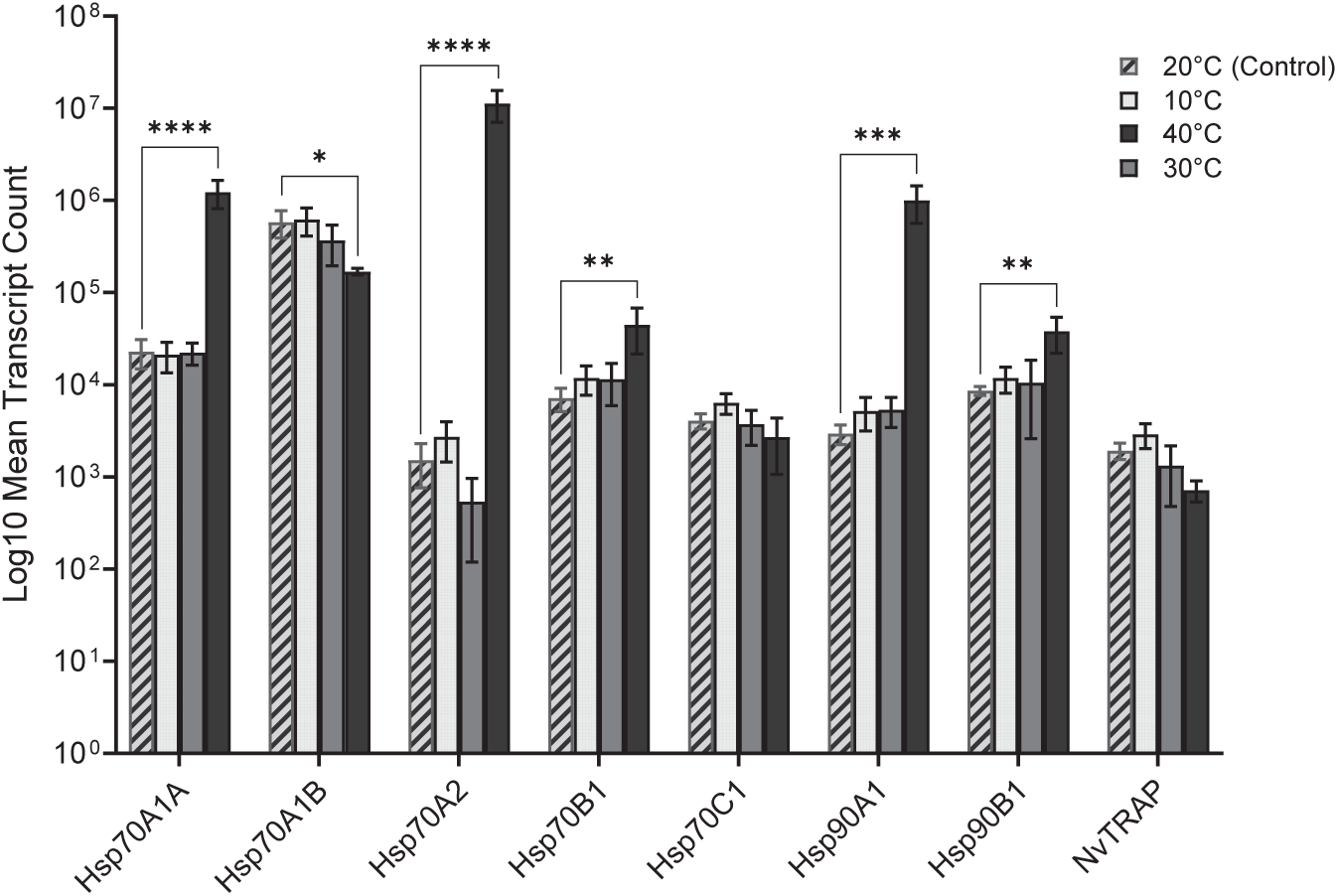
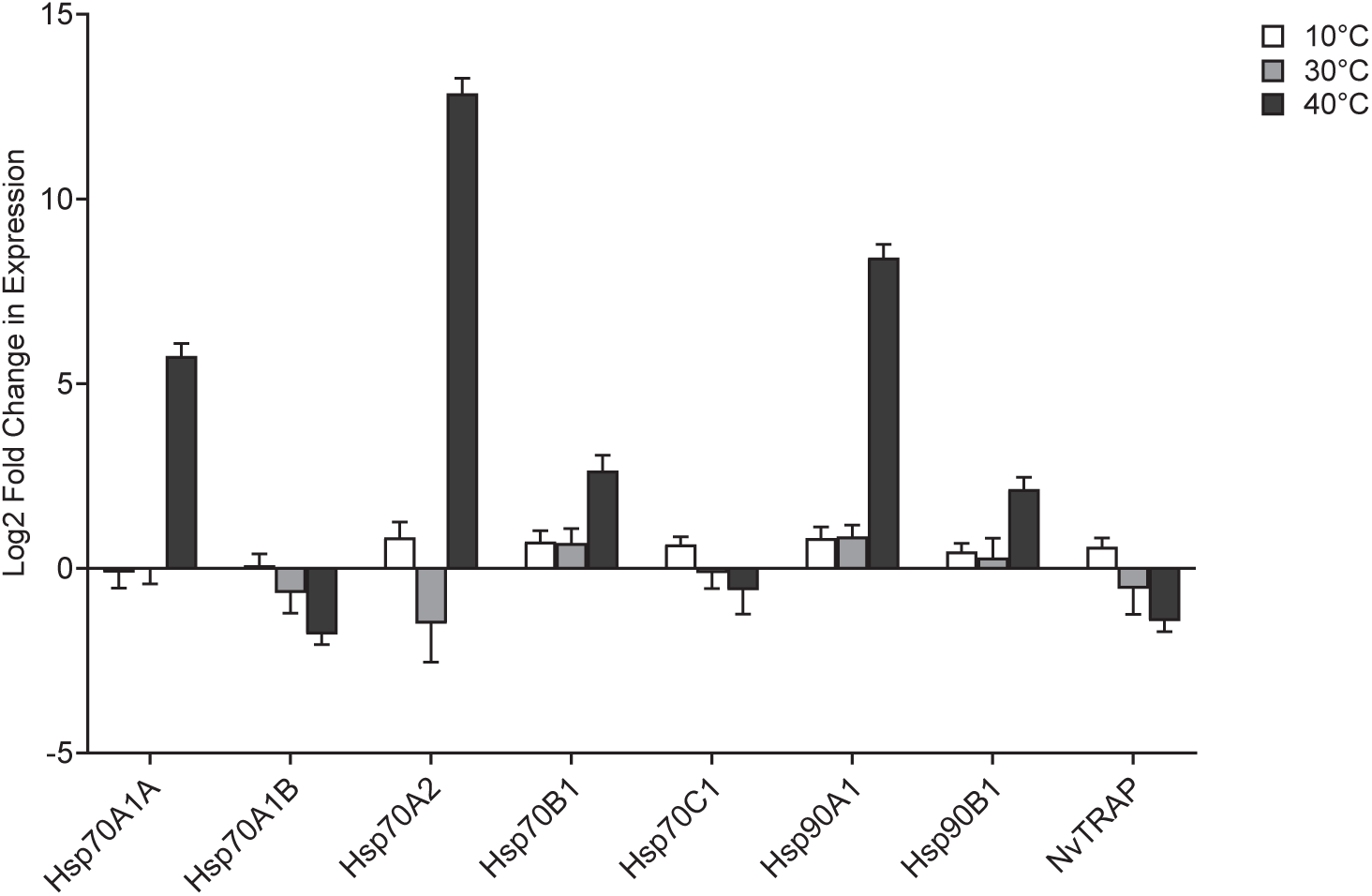
Quantitative PCR data from *N. vectensis* anemones across temperature treatments. A: Bar plot of HSP transcript abundance across three experimental temperatures and one control temperature. Plotted values are averages of four replicates for 10°C and 20°C, and three replicates for 30°C and 40°C conditions, respectively. Error bars represent the standard error. Asterisks denote significance calculated with a one-way ANOVA test followed by a Dunnett’s test. P-values are denoted by * for p≤0.05, ** for p≤0.005, *** for p≤0.0005, and **** for p≤0.0001. B: Bar plot of HSP fold change in expression across three experimental temperatures. Plotted values are replicate means normalized to the mean of control (20°C) with a log 2- transformation of the fold change values for better visualization. Error bars represent standard error of the mean.

### 2.4. Expression of HSP70s and HSP90s over *N. vectensis* development

We compared the expression of each HSP gene across five developmental stages - embryo, early larva, late larva, juvenile, and adult. We found Hsp70A1B as the sole HSP to have an average increase in expression between all the developmental stages studied (Figure 4A). In addition, Hsp70A1B displayed the highest mean expression among all HSPs in all stages except the embryo (Figure 4A). In contrast, Hsp70A2 showed the lowest mean expression in all stages except the juvenile stage (Figure 4A). All HSPs showed a trend of higher expression in the early larval stage compared to the embryo stage, with significantly higher expression of Hsp70A1B, Hsp70A2, Hsp70B1, and Hsp90B1 (p<0.05, Figures 4B and 4C). Overall, Hsp70A1A, Hsp70B1, Hsp90A1, Hsp90B1, and NvTRAP showed a similar expression pattern with a peak in expression during the early larval stage, followed by lower expression during the juvenile stage (Figure 4A).

**Figure 4.**
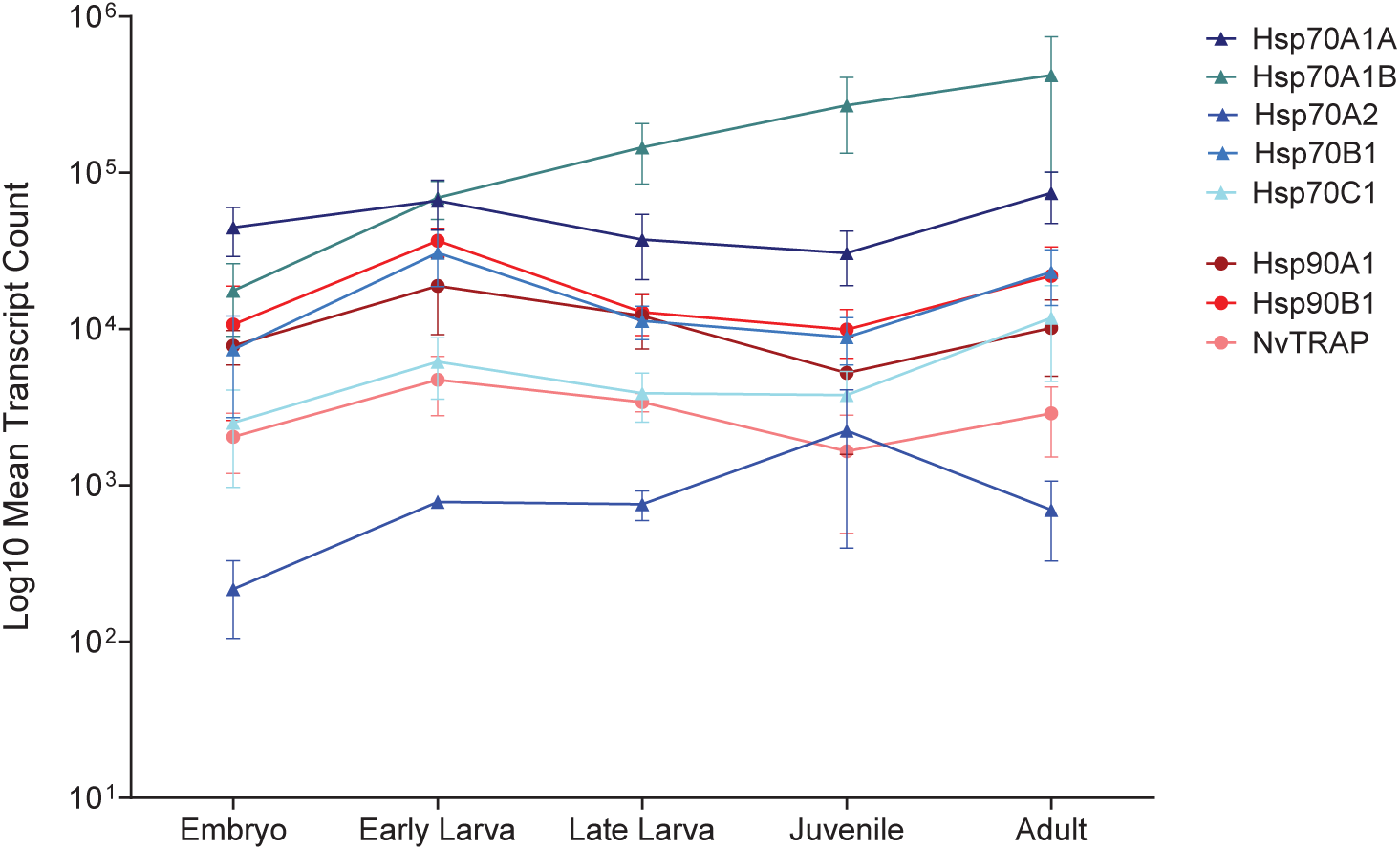
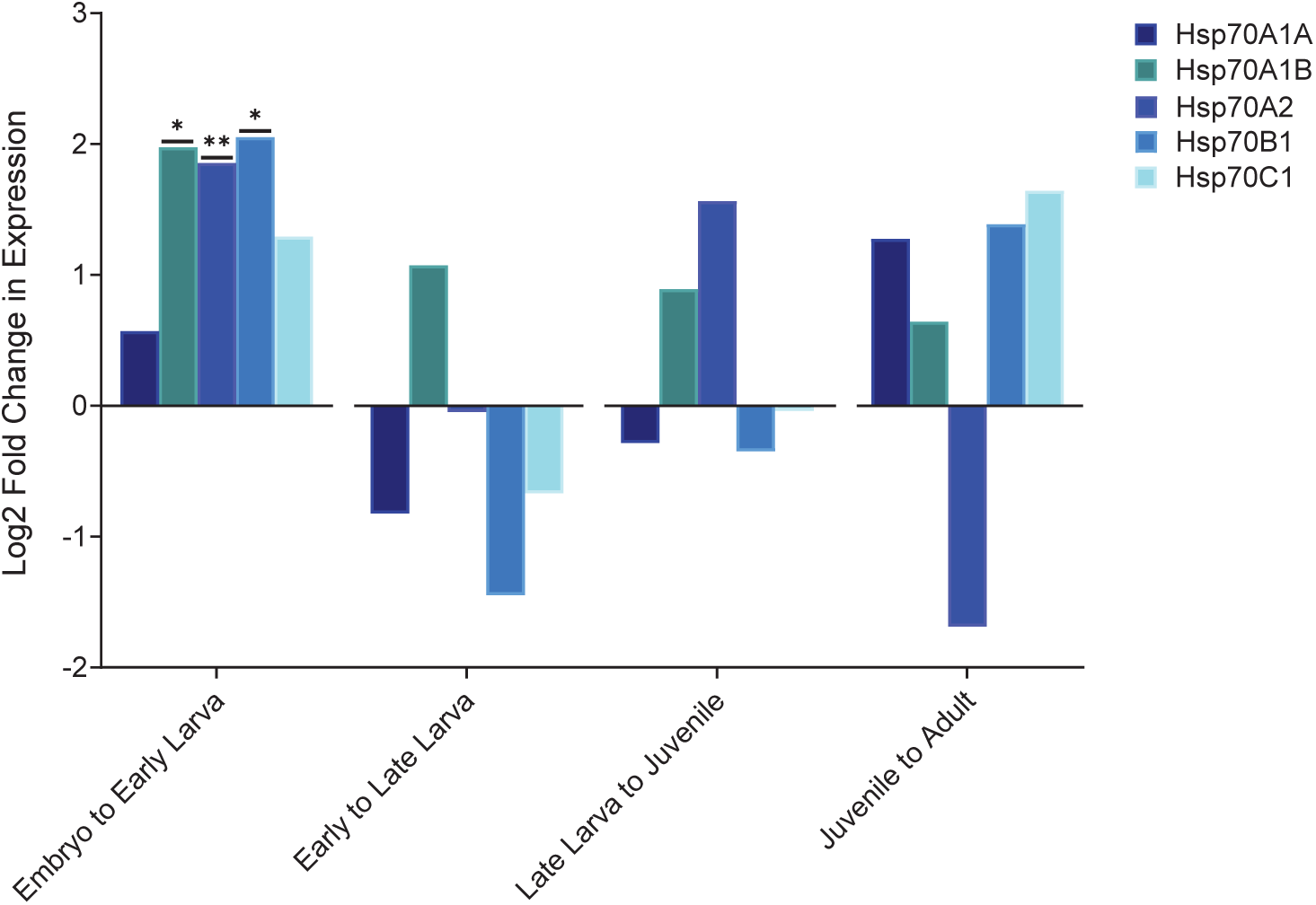
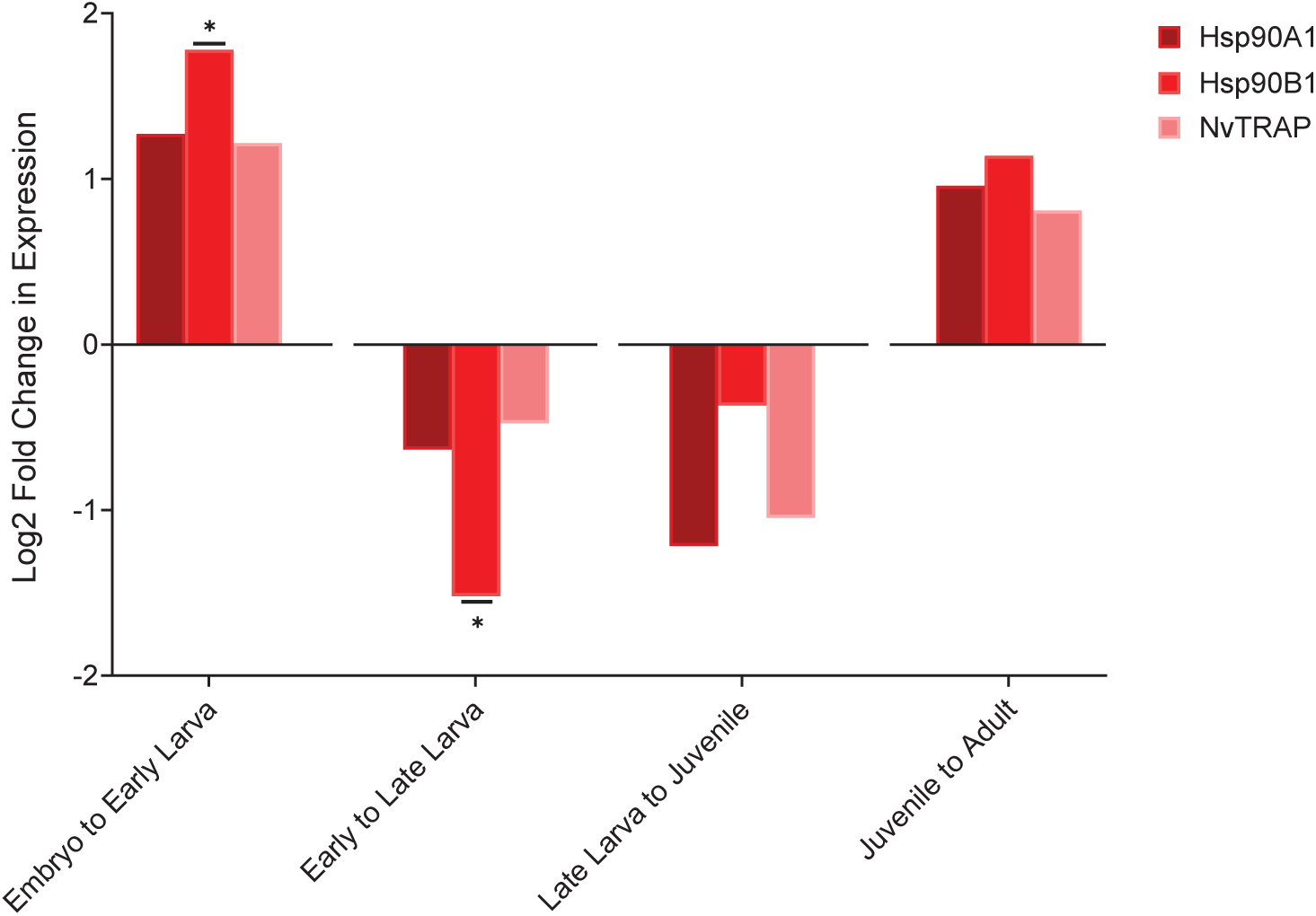
Quantitative PCR data from *N. vectensis* anemones across developmental stages. A: Line graph of HSP transcript abundance across five developmental stages. Plotted values are averages of four replicates for each HSP and stage combination. Error bars represent the standard deviation. B: Bar plot of HSP70 fold change in expression between consecutive developmental stages. Plotted values are log2- transformed fold changes between the expression at each developmental stage with respect to the previous developmental stage. Asterisks denote significance calculated with a two-sample two-tailed t-test and Benjamini FDR-corrected p-values. C: Bar plot of HSP90 fold change in expression between consecutive developmental stages. Plotted values are log2-transformed fold changes between the expression at each developmental stage with respect to the previous developmental stage. Asterisks denote significance calculated with a two-sample two-tailed t-test and Benjamini FDR-corrected p-values. P-values are denoted by * for p≤0.05 and ** for p≤0.005.

### 2.5. Cell-type specific expression of *N. vectensis* HSPs during development

We leveraged the published developmental single-cell atlas by Steger, Cole [39] to compare the expression of *N. vectensis* HSPs in different cell types of this species. Overall, HSPs showed broad expression patterns across 12 cell types (Figure 5A). Notably, a high proportion of cells from all 12 clusters expressed Hsp90A1 and Hsp70A1A, with the highest average expression in the neuronal cluster. Most clusters also had a considerable number of cells expressing high levels of Hsp70A1B, with the exceptions of cnidocyte and embryonic endomesoderm clusters. In comparison, Hsp70A2 had moderate and Hsp90B1, Hsp70B1, and Hsp70C1 had low expression levels across clusters. Hsp90C1 was excluded from this analysis on account of poor expression across all clusters (data not shown). The broadly expressed cytosolic Hsp70A1A and Hsp90A1 exhibited distinctly similar expression patterns as confirmed by principal component analysis (PCA) (Figures 5A and 5B). Likewise, endoplasmic reticulum-associated Hsp70B1 and Hsp90B1 depicted similar expression patterns with high expression in the cnidocytes, but not in the mature cnidocytes. In contrast, Hsp70A2 and Hsp70A1B were highly expressed in mature cnidocytes, but not in the general cnidocyte cluster.

**Figure 5.**
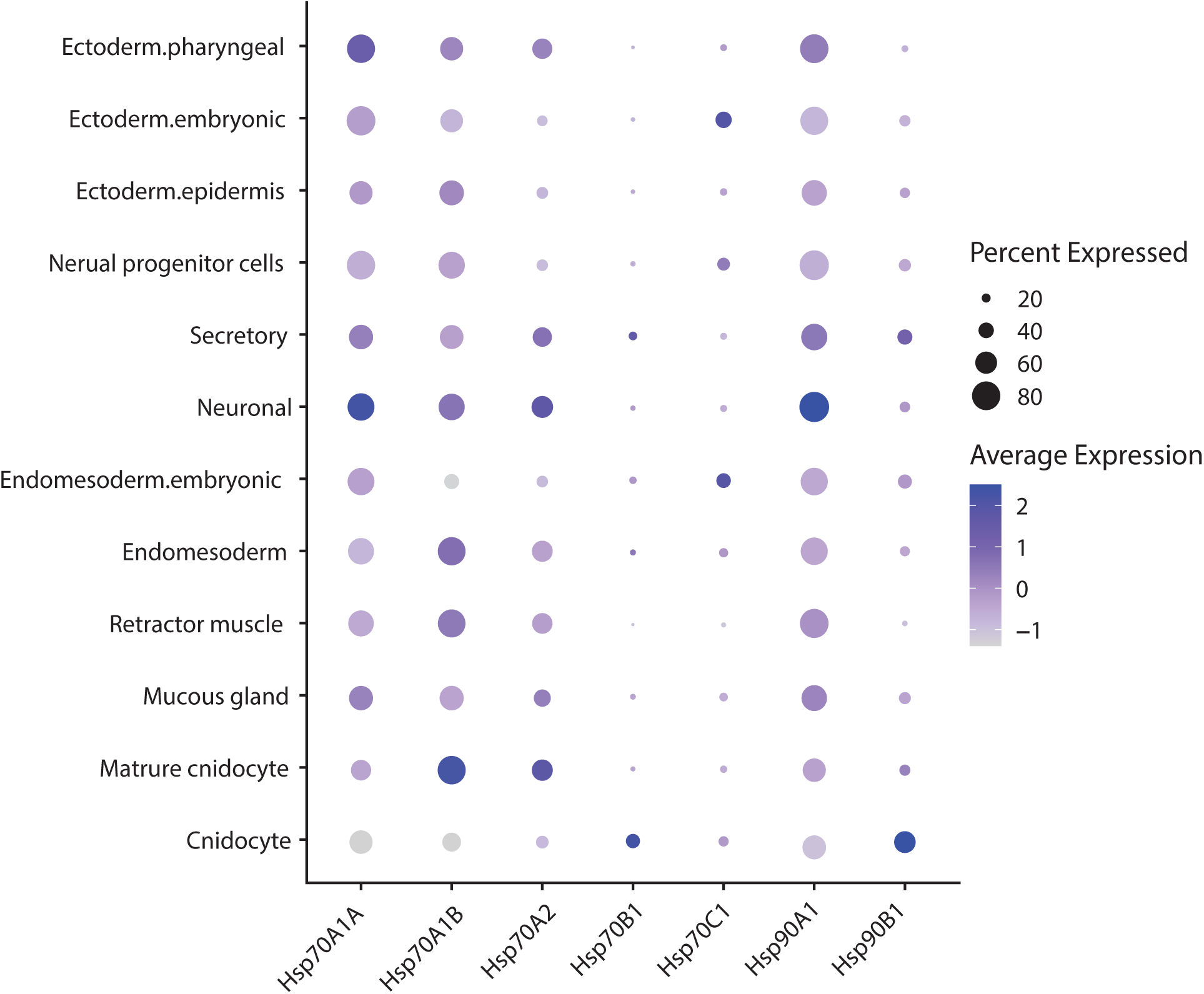
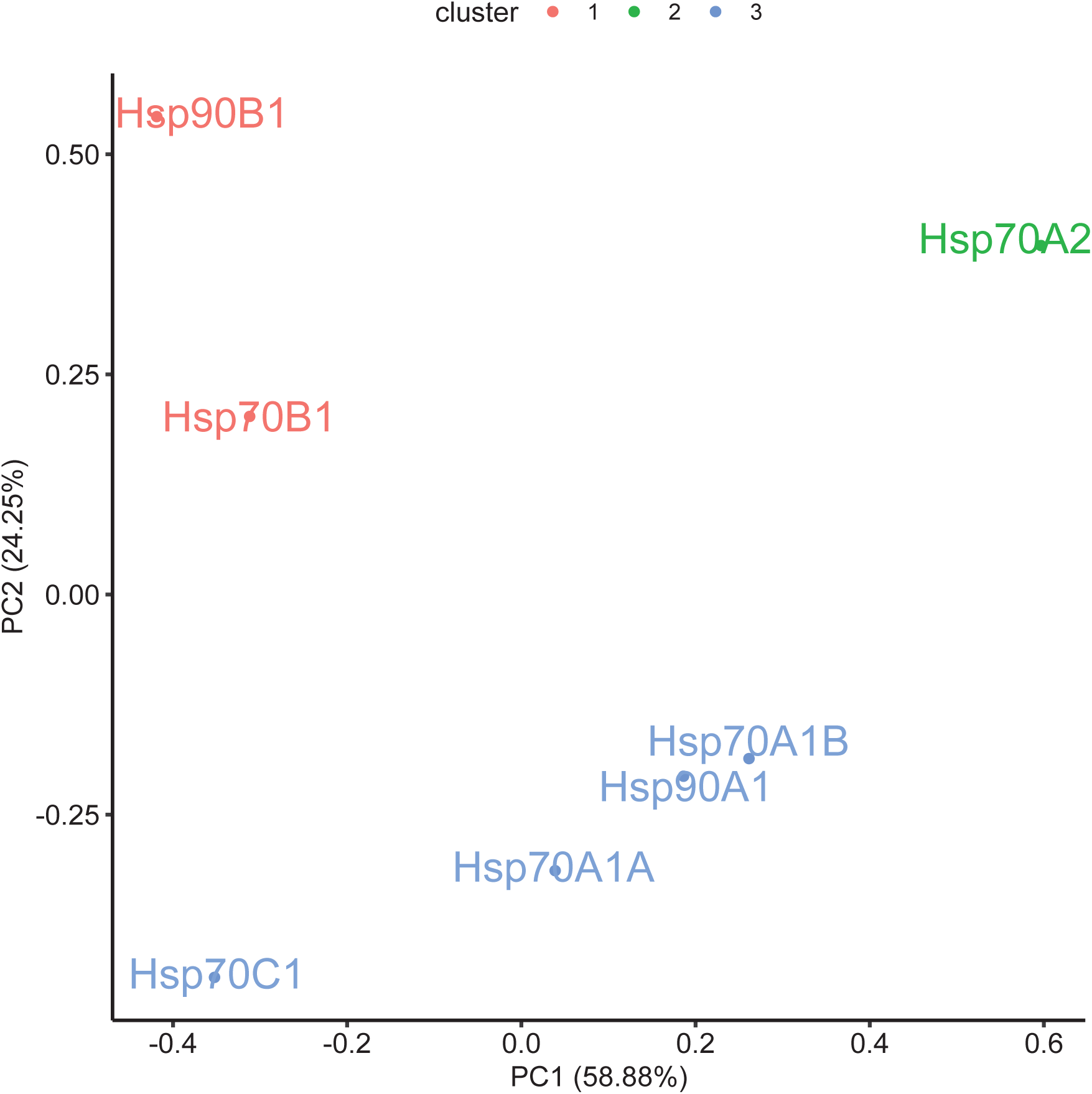
Single-cell transcriptomic data from *N. vectensis* anemones. A: Dot plot of HSP expression levels across 12 cellular clusters. The size of each dot represents the percent of cells in each cluster expressing the corresponding gene, while the dot color represents the average log-normalized expression level across the cluster. B: Principal component analysis (PCA) of log-normalized expression levels of HSPs obtained from the single-cell RNA sequencing data. Three computationally confirmed clusters are marked in different colors (red, blue, and green), denoting groups of HSPs with similar expression levels.

### 2.6. Distribution of *cis*-regulatory heat shock elements in HSP promoters

To identify potential differences in the regulation of each HSP gene, we identified the subunit number, orientation, and type of HSE motifs within 500 bp upstream sequences of *N. vectensis* HSP transcription start sites (TSS). We identified a total of nine HSE motifs in the promoters for four HSP genes (Figure 6). All identified HSEs were located within 350 bp of the TSS. Most of the identified HSEs were comprised of three alternating pentameric subunits of ‘nGAAn’ and ‘nTTCn’ repeats; however, one motif each in Hsp70A1A and Hsp70A2 had four pentameric subunits, and one motif in Hsp90A1 had six subunits. Of the nine total HSEs identified, five were head-to-head (‘nGAAn’ is the most distal subunit from TSS) and four were tail-to-tail (‘nTTCn’ is the most distal subunit from TSS) motifs. Interestingly, all motifs located in the HSP90 promoters were tail-to-tail type motifs, while only one motif, the left-most motif for Hsp70A2, was in the tail-to-tail orientation.

**Figure 6.**
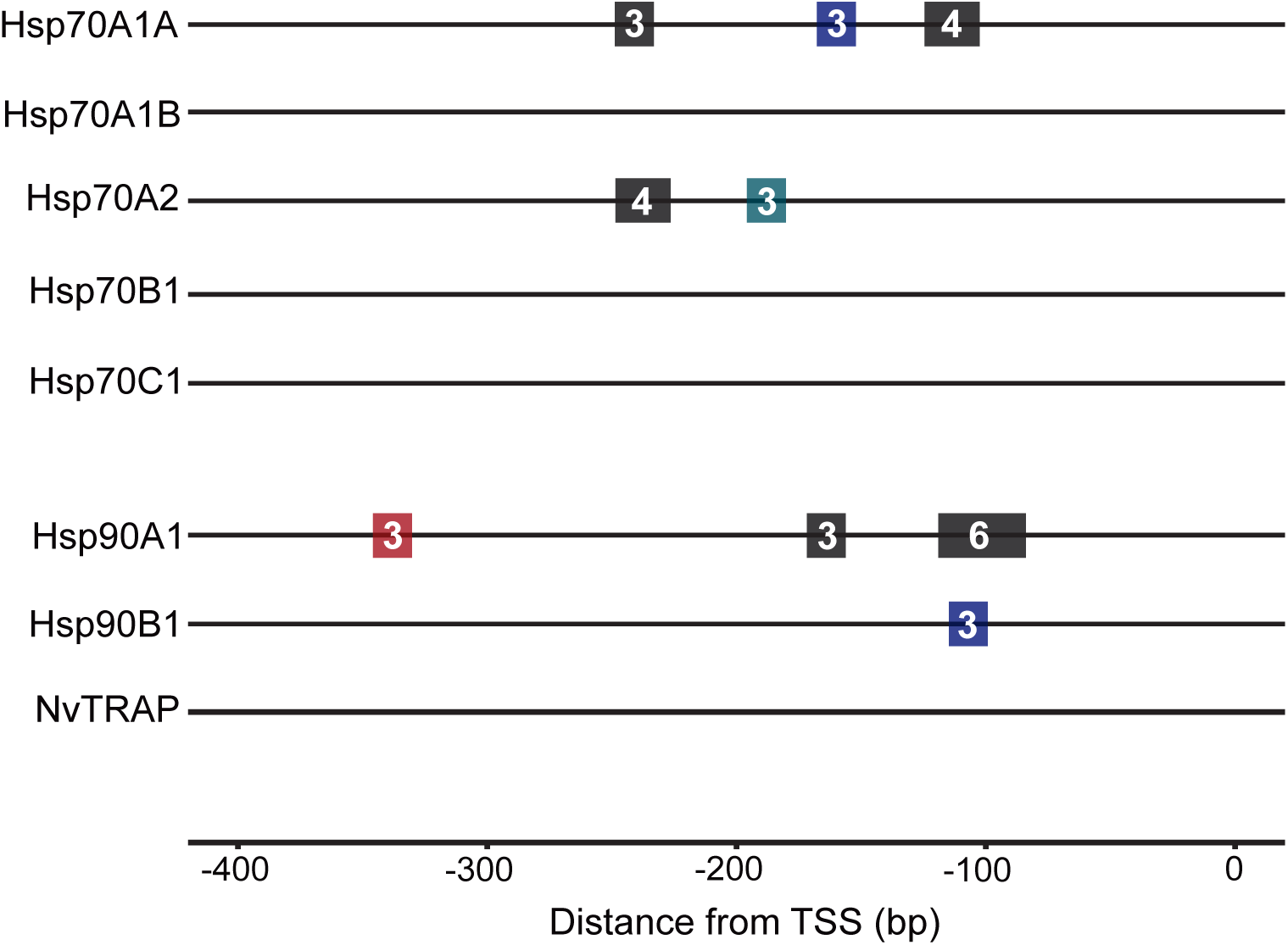
Promoter heat shock element architecture of HSP70 and HSP90 genes from *N. vectensis.* Horizontal lines represent the promoters, and the transcription start site (TSS) is positioned at zero (right end) for each promoter. Rectangles represent HSE motifs (for better visualization, motifs are illustrated 30% longer than their actual length). The numbers within rectangles represent the number of individual ‘nGAAn’ and ‘nTTCn’ pentamers within the motif. Varied type motifs are denoted in black, gapped type in blue, typical type in green, and key type in red.

We further characterized HSEs as being typical (having the canonical HSE sequence), gapped (having a mismatched base in an interior subunit), varied (having a mismatched base in an exterior subunit, except in the key positions of ‘G’ in ‘nGAAn’ or ‘C’ in ‘nTTCn’), or key (having a mismatched base in an exterior subunit in one the key positions of ‘G’ or ‘C’) type motif. Overall, we found seven varied HSE motifs, of which three were in the Hsp70A1A promoter, two were in the Hsp90A1 promoter, and one each was in the Hsp70A2 and Hsp90B1 promoters. Intriguingly, we only detected one typical HSE, and it was in the Hsp70A2 promoter. There were no gapped HSEs present in the promoters examined; however, we found a key HSE in the Hsp90A1 promoter. The promoters of Hsp70A1B, Hsp70B1, Hsp70C1, and NvTRAP had no detected HSE motifs. Hsp70A1A and Hsp90A1 had the highest number of HSEs in their promoters, and these were also highly induced under heat stress (Figures 3A and 6).

## 3. Discussion

Several studies have provided evidence for lineage-specific variation for expression, induction, and regulation of HSPs across diverse ecological niches [14–18]. In *Nematostella vectensis*, prior work showed that three cytosolic members of the HSP70 family exhibit distinct induction patterns under heat stress and differ in their ability to protect cells from stress and in protein-protein interactions as chaperones [34, 38]. Building on this research, our study characterizes the phylogenetic relationships, heat stress induction, developmental expression, and regulatory motifs of all HSP70 and HSP90 family members in *N. vectensis*.

Characterizing the number and diversity of HSP70s and HSP90s is a prerequisite to comprehensively compare shared and unique HSP functions for a species. While HSP expression for *N. vectensis* has been reported in a number of previous studies [31, 33, 35, 37], the identity of these genes and how they represent the full complement of HSPs remained unclear. Here, we identified five distinct HSP70 and three HSP90 genes in *N. vectensis*. A sixth HSP70 gene appeared to be either a very recent duplication or an artifact of genome assembly due to the identical coding and promoter sequence and was not included in the analyses. The identified HSP70s and HSP90s from *N. vectensis* are generally representative of other eukaryotic species with at least one homolog in each phylogenetic grouping that corresponds to subcellular localization (cytoplasm, endoplasmic reticulum, and mitochondria). In other metazoan taxa, HSPs belonging to various localization-based clades show distinct functions [40]. Like many other species, *N. vectensis* has multiple HSP70s in the cytoplasm-nuclear clade. Hsp70A2 resulted from a deep evolutionary split, likely at the origin of the animal kingdom, while Hsp70A1A and Hsp70A1B appear to have resulted from a recent duplication event. Further phylogenetic analyses with additional cnidarian species will be informative to better characterize the timing of this gene duplication event.

We exposed *N. vectensis* to a range of environmentally relevant experimental temperatures: 10°C, 20°C, 30°C, and 40°C for 6 hours to mimic the breadth of the conditions experienced in the wild. Consistent with the heat stress expression data of Waller, Knighton [34], our results show that a 10°C acute increase or decrease did not significantly induce any of the HSPs studied, suggesting a higher thermal induction threshold compared to other marine invertebrates. By contrast, in corals and other sea anemones, HSP70 gene expression and protein production have been induced by smaller temperature increases (4–10 °C) within the first 0–4 hours of stress exposure [19, 41–43]. We hypothesize that the elevated threshold in *N. vectensis* likely reflects physiological adaptation to its extreme and variable estuarine habitat, where daily temperature fluctuations of 15–20 °C are common [27]. These findings support the hypothesis that HSP expression thresholds are evolutionarily tuned to match an organism’s thermal niche and that inducibility varies across orthologs or taxa. In *N. vectensis*, substantial induction of transcription was observed in Hsp70A1A, Hsp70A2, and Hsp90A1 following a 20°C increase, confirming them as the inducible cytosolic HSPs. Among these, Hsp70A2 showed the strongest induction, consistent with previous findings that yeast expressing this gene as their sole HSP70 exhibit high resistance to diverse stressors [34]. By contrast, Hsp70A1B was constitutively expressed at all temperatures, in agreement with earlier observations in *N. vectensis* [34].

HSPs, particularly HSP90, have critical roles in the development of diverse species [44]. In *Drosophila melanogaster,* HSP90 is critical for successful development of nearly all adult structures due to its role in maintaining the activity of signal transduction proteins in a variety of developmental pathways, regulating processes such as cell cycle, apoptosis, and differentiation [45, 46]. We examined HSP expression across five developmental stages in *N. vectensis*. Overall, HSPs were dynamically expressed throughout development, with each HSP displaying both unique and overlapping patterns. Six of the eight HSPs showed highly similar expression profiles, characterized by elevated expression during the early larval stage and decrease at the juvenile stage. This pattern suggests that HSP genes may play both shared and distinct developmental roles, raising the question of whether they function redundantly or are individually essential for proper development in marine invertebrates. Notably, the stress-inducible Hsp70A2 exhibited the lowest expression across stages, suggesting a primary function in stress response. In contrast, its stress-inducible partner Hsp70A1A displayed the second-highest expression across stages, while the constitutively expressed Hsp70A1B showed the highest expression overall. These patterns suggest that Hsp70A1A and Hsp70A1B may have broader functions, contributing to both stress response and developmental processes. Further research to determine spatial expression and manipulate gene expression would be informative in understanding the potential roles of these HSPs in development.

Single-cell transcriptomic data revealed cell-type-specific expression patterns during development, suggesting cell-specific roles for distinct HSPs, particularly in neuronal and cnidocyte cell populations. Cnidocytes are specialized stinging cells unique to cnidarians that function in prey capture and defense, and prior studies have shown that stress response genes are expressed in this cell type. For instance, in adult *N. vectensis*, the generalized stress response transcription factor NF-KB was shown to be required for cnidocyte development [47, 48]. Similarly, in the coral *Acropora hyacinthus*, stress response genes such as *AP-1*, *FosB*, and *TNFR41* were expressed in cnidocytes under heat stress, suggesting a role for this cell type in mediating stress responses [49]. Our findings extend these observations by also suggesting that HSPs may contribute to cnidocyte development or perhaps be important in maintaining cnidocytes by preventing proteotoxic conditions in that cell type. Further research will be necessary to determine whether HSPs are specifically deployed in cnidocytes as a part of stress response processes.

Finally, we identified differences in both the number and type of HSEs present in the promoters of *N. vectensis* HSP genes. The highest number of HSEs were found in the promoters of strongly stress-induced genes (Hsp70A1A, Hsp70A2, and Hsp90A1), consistent with other species, where greater numbers of promoter HSEs correlated with stronger induction under stress [50]. In contrast, HSPs lacking HSEs in their promoters are likely regulated through mechanisms outside the canonical HSF regulation. We also observed variation in HSE type, with some promoters containing canonical motifs and others containing mismatched versions. For most HSPs with motifs, it was difficult to correlate HSE sequence variation with stress-induced expression because of the presence of multiple HSE types within the same putative promoter. Hsp90B1 was the only gene with a single HSE in its promoter which we identified as being a gapped type of motif. A potential reason for the lack of heat stress-induced expression of Hsp90B1 could be that the sequence of the gapped type HSE found in its promoter might not allow for efficient HSF binding. This, however, stands in contrast to previous findings in bread wheat, where it was reported that gapped and varied type sequence variations showed comparable heat stress response magnitude to typical type HSEs [51]. Further research will be needed to determine whether HSE type influences HSP expression profiles in *N. vectensis*. Overall, these promoter differences provide a potential mechanistic basis for the diverse expression patterns observed, supporting the view that cis-regulatory evolution has contributed to functional diversification within the HSP gene family.

Given the shared and unique expression patterns of HSPs during heat stress and development in *N. vectensis*, we suggest caution when using the general terms ‘HSP70’ or ‘HSP90’ as reliable universal biomarkers of stress. Although HSP induction has long served as a molecular indicator of proteotoxic stress [1], our data, as well as previous data in other species, shows that phylogenetic relationships of genes in these chaperone families is important. Inducible and cognate types of HSP genes have evolved repeatedly in eukaryotes, thus precise comparisons of evolutionary relationships and expression are required to determine specific biomarkers. For *N. vectensis*, some HSPs are upregulated only under extreme conditions, while others are highly expressed during development in the absence of external stressors. While a growing body of evidence highlights the diverse roles of HSPs in model organisms, a comprehensive functional characterization of cnidarian HSP family genes is still lacking [19]. Future work should therefore focus on identifying HSP orthologs that are most consistently and specifically expressed under stress across diverse cnidarian species, thereby narrowing down candidates that may serve as reliable biomarkers of generalized stress and other functions.

## 4. Materials and Methods

### 4.1. Culturing of animals

Adult *Nematostella vectensis* originally collected from the Great Sippewissett Marsh, Massachusetts, were cultured under standard conditions (room temperature, 13 parts per thousand (ppt) artificial seawater (ASW), fed freshly hatched *Artemia nauplii* 2-3 times per week, and weekly water changes).

To procure developmental stages, we used a standard weekly feeding with mussel tissues followed by an overnight temperature increase of 25°C with full lighting. After spawning, egg masses containing embryos were isolated from the bowl containing adults, the jelly was removed with a 4% cysteine wash, and then separately cultured in 13 ppt seawater. We isolated individuals from multiple egg masses at the embryo (1 days post fertilization (dfp)), early larva (3-4 dpf), late larva (7-11 dpf, not metamorphosed), and juvenile stages (8-23 dpf, metamorphosed), removed excess seawater, and then snap froze with liquid nitrogen until extraction of RNA.

We performed an acute temperature change experiment with adult anemones to determine how the transcription of the identified HSP genes was influenced by abrupt temperature changes. Adult anemones were cultured in the dark at 20°C for more than one month before temperature exposure. We separated individual anemones into glass bowls (n=4 replicate bowls per temperature) with filter-sterilized ASW that had been equilibrated to four temperatures: 10°C, 20°C, 30°C, and 40°C. Anemones were then incubated at the set temperature for 6 hours. The anemones were removed from the bowl with a pipette and placed in individual 1.5 ml microcentrifuge tubes. Excess seawater was removed, and the samples were snap-frozen in liquid nitrogen and stored at -80°C until RNA extraction.

### 4.2. Identification of heat shock proteins

HSP70 and HSP90 genes from *N. vectensis* were identified through a combination of BLASTp and BLASTx searches of the proteome and genome assemblies, through the Joint Genome Institute and SIMRbase, respectively. Subcellular localization of HSPs was predicted using the high-throughput model of DeepLoc 2.1 [52]. We used HSP70 and HSP90 protein sequences from each clade of human and *Drosophila melanogaster* types of HSP family corresponding to subcellular locations (cytoplasmic-nuclear, endoplasmic reticulum, mitochondria) as query sequences. We compared each retained BLAST hit (e^-10^) with an alignment to remove duplicate annotations in the genome. We confirmed the predicted sequence for each gene through amplification, cloning and Sanger sequencing of portions of each predicted HSP transcript [primers designed with Primer3 [53], primer sequences listed in Table S1]. PCR products were gel purified, ligated into pGEM-T Easy vector (Promega), and sequenced to confirm the targeted amplicon. These cloned sequences were also used for generation of a standard curve for quantification of gene expression with RT-qPCR (see section 4.4).

### 4.3. Phylogenetic analysis

We used maximum likelihood to determine the evolutionary relationships of the identified HSP70 and HSP90 proteins for *N. vectensis* with selected animal, fungal, and bacterial species. HSP70 sequences were selected from Boorstein, Ziegelhoffer and Craig [54] that previously described the diversity of HSP70 genes in a diversity of eukaryotes. Similarly, we selected from sequences used in a study of the phylogenetic diversity of HSP90 genes in eukaryotes by Chen, Zhong and Monteiro [55]. We included sequences for these species from each of the subcellular localizations to characterize the diversity of these chaperones in *N. vectensis*. We also included HSP70 sequences from the coral *Acropora millepora* [56] and *Hydra magnipapillata* [57] identified through BLAST to better resolve relationships in this family because a unique clade of HSP70 genes has been reported in non-bilaterian animals including cnidarians. [40]. Full length sequences for all taxa were aligned with Muscle 3.6 [58] and the ends containing unalignable regions were removed. Maximum likelihood analyses were run using RAxML (version 7.0.4) [59] with a RtREV+G matrix (model determined by AIC criteria with ProtTest v1.4). Trees were visualized and illustrated with FigTree v1.4.4 available at GitHub. For gene naming, we used the nomenclature corresponding to subcellular locations for HSP70s: “A” for cytoplasmic-nuclear, “B” for endoplasmic reticulum, and “C” for mitochondria. HSP90 genes were named after the nomenclature by Chen, Zhong and Monteiro [55]: “A” for cytoplasmic-nuclear, “B” for endoplasmic reticulum, “C” for plastids (e.g, chloroplast) and “TRAP” for mitochondria.

### 4.4. RNA extraction and quantitative real-time PCR

For characterization of developmental expression and temperature response patterns, RNA was extracted using the Aurum Total RNA Mini Kit with on-column DNAse digestion. cDNA was synthesized from total RNA with the Iscript cDNA Synthesis Kit using 2 µg of RNA per 30 µl reaction. For quantitative PCR, we followed the approaches previously reported in Reitzel and Tarrant [60]. Briefly, the PCR cycle conditions were: 95°C for 3 min; 40 cycles of 95°C for 15 s and 64°C for 45 s. Oligonucleotide primers (see Table S1) were designed to amplify each *N. vectensis* HSP gene. Primers were 20 nt, with a GC content of 40-60%, either overlapped predicted exon-exon boundaries by 3-4 bp or spanned a large intron (where possible), and produced predicted amplicons of <150 bp with minimal predicted secondary structure [m-fold, [61]]. A standard curve was constructed from serially-diluted plasmids containing the amplicon of interest. The standard curve was used in qPCR reactions to calculate the number of molecules per reaction. qPCR was performed using iQ SYBR Green Supermix (Bio-Rad), and reactions were run a MyCycler Real-Time PCR detection system (Bio-Rad). After the completion of the amplification cycles, the PCR products were subjected to melt curve analysis to ensure that only a single product was amplified. The number of molecules per µl for each gene was calculated by comparing the threshold cycle (Ct) from the sample with the standard curve. Expression was compared between developmental stages or temperatures using one-way analysis of variance (ANOVA) with Tukey’s Honestly Significant Difference Test as a post hoc test.

### 4.5. Cell type-specific expression analysis

To gain further insights into the expression of HSPs at the cellular level, we interrogated scRNA-seq data across a time course of *N. vectensis* developmental stages published by Steger, Cole [39]. Counts matrix and cell barcodes were downloaded from the Gene Expression Omnibus (GEO: GSE200198), and analysis was performed using previously published methods (https://github.com/technau/CellReports2022/blob/main/DataS6_Merge2Figures.R). The fold change for each HSP was extracted for each cell cluster with a p-value of <0.01 and subsequently used as input for principal component analysis, performed in R.

### 4.6. Identification and distribution of regulatory elements

The 500 bp putative promoter sequences of candidate HSP70s and HSP90s were extracted from the *N. vectensis* genome using a previously developed python script [43]. These sequences were searched for the canonical heat shock element (HSE) sequence ‘GAANNTTCNNGAA’ using the FIMO tool from MEME Suite (https://meme-suite.org/meme/tools/fimo). Identified motifs were manually examined for overlap and subunit numbers were recorded for each unique HSE location. Unique motifs were characterized as head-to-head (‘nGAAn’ is the most distal subunit from TSS) or tail-to-tail (‘nTTCn’ is the most distal subunit from TSS) motifs as previously described [62, 63]. Further characterization involved identifying each motif as being typical (having the canonical HSE sequence), gapped (having a mismatched base in an interior subunit), varied (having a mismatched base in an exterior subunit, except in the key positions of ‘G’ in ‘nGAAn’ or ‘C’ in ‘nTTCn’), or key (having a mismatched base in an exterior subunit in one the key positions of ‘G’ or ‘C’) type motif based on methods described in Zhao, Javed [51]. The ‘ggplot2’ package was used in R to visualize the promoter HSE architecture for all HSPs.

## Acknowledgements

This research was supported by the National Science Foundation grant number 1545539 to AMR. We would like to thank Ann M. Tarrant for helping with conceptualization of this research project, provision of materials and resources, and reviewing the manuscript draft.

## Author Contribution

JAB: validation, formal analysis, visualization, writing – original draft, writing – reviewing and editing. JMS: formal analysis, visualization, writing – reviewing and editing. AMR: conceptualization, validation, investigation, supervision, funding acquisition, writing – reviewing and editing.

## Declaration of Interest

The authors declare no competing interests.

**Abbreviations**
HSP: heat shock protein;
ER: endoplasmic reticulum;
HSF: heat shock factor;
HSE: heat shock element;
ANOVA: analysis of variance;
qPCR: quantitative real-time polymerase chain reaction;
FDR: false discovery rate;
PCA: principal component analysis;
TSS: transcription start site;
NF-KB: nuclear factor kappa-light-chain-enhancer of activated B cells;
AP-1: activator protein 1;
FosB: FBJ murine osteosarcoma viral oncogene homolog B;
TNFR41: tumor necrosis factor receptor 41;
PPT: parts per thousand;
ASW: artificial seawater;
DPF: days post fertilization;
RNA: ribonucleic acid;
DNA: deoxyribonucleic acid

