## Supplementary Material for "Transcription dynamics and regulation of heat shock protein genes during stress and development in the estuarine cnidarian *Nematostella vectensis*"

| Gene name | Cloned Fragment Primers (5'-3') | Product Size (bp) | qPCR Primers (5'-3') | Product Size (bp) |
|---|---|---|---|---|
| HSP70A1A | TTTTCCAACACGGAAAGGTC | 818 | AATGAACCCCGAGAACACAG | 52 |
| | ATCAATGCCCTCAAACAAGG | | TGGGTCATATCACGAGCAAC | |
| HSP70A1B | CAAACGACCAAGGAAATCGT | 1001 | TCGATGATCCTGGGGTAAAG | 74 |
| | GGACTTGTTCAGCTCCTTGC | | CCTGCCTCGTTCACTACCTC | |
| HSP70A2 | CTTTTGCTGCCGAAGAAATC | 972 | TCATCCAGCACTGAAGCAAG | 72 |
| | GCTGGTTGTCGGAATACGTT | | CTCGGCTGATTTTCGTGTAG | |
| HSP70B1 | ATTATCGCAAACGACCAAGG | 939 | TGAGATGCTCACTCGTGCTC | 78 |
| | AGAGCCTCCGACAAGAACAA | | ACTTTCTGGACGGGCTTCAG | |
| HSP70C1 | TTCAGTGGTTGCCTTTACCC | 1000 | ACACCCTATACGCCACCAAG | 127 |
| | AGCACCGATAGCAACAGCTT | | CAACTTCAAACCAAGCATCG | |
| HSP90A | TCGCTCAGTTGATGAGCTTG | 959 | ACCATTGCCAAGAGTGGAAC | 90 |
| | GTCGAAAGGTGCTCTCTTGG | | GAAACCCACACCGAATTGAC | |
| HSP90B | CCTGCACTTGAAGGAAGAGG | 1111 | CGAGGTTGAAGAGGACAAGG | 80 |
| | CTTGTCGCTGTCTTCACCAA | | GTTCCCAGTCCCACACAGTC | |
| HSP90TRAP | GACCGGACCTTCTAACACCA | 1073 | GTCAATCAATGCGAGTGTGC | 96 |
| | TCGGGGTAACACCTTTTCAG | | GGTTCTTCCACATCCTCACC | |

**Table S1.** Primer sequences used for cloning and qPCR.
